# Insect wings and body wall evolved from ancient leg segments

**DOI:** 10.1101/244541

**Authors:** Heather S. Bruce, Nipam H. Patel

## Abstract

The origin of insect wings has long been debated. Central to this debate is whether wings evolved from an epipod (outgrowth, e.g., a gill) on ancestral crustacean leg segments, or represent a novel outgrowth from the dorsal body wall that co-opted some of the genes used to pattern the epipods. To determine whether wings can be traced to ancestral, pre-insect structures, or arose by co-option, comparisons are necessary between insects and arthropods more representative of the ancestral state, where the hypothesized proximal leg region is not fused to the body wall. To do so, we examined the function of five leg patterning genes in the crustacean Parhyale hawaiensis and compared this to previous functional data from insects. By comparing gene knockout phenotypes of leg patterning genes in a crustacean with those of insects, we show that two ancestral crustacean leg segments were incorporated into the insect body, moving the leg’s epipod dorsally, up onto the back to form insect wings. Thus, our data shows that much of the body wall of insects, including the entire wing, is derived from these two ancestral proximal leg segments. This model explains all observations in favor of either the body wall origin or proximal leg origin of insect wings. Thus, our results show that insect wings are not novel structures, but instead evolved from existing, ancestral structures.

**One Sentence Summary:** CRISPR-Cas9 knockout of leg gap genes in a crustacean reveals that insect wings are not novel structures, they evolved from crustacean leg segments

## Main Text

The origin of insect wings has fascinated researchers for over 130 years. One theory proposes that the proximal portion of the ancestral crustacean leg became incorporated into the body (*1*), which moved the leg’s epipod (lobe-shaped outgrowth, e.g. gill) dorsally, up onto the back to form insect wings (*2*). Another theory proposes that the wing is a novel outgrowth from the dorsal body wall that co-opted some of the genes used to pattern the epipods of leg segments (*3*). Alternatively, wings may be derived from a combination of leg and body wall (dual origin, (*4*)). To determine whether wings can be traced to ancestral, pre-insect structures, or arose by co-option, comparisons are necessary between insects and other arthropods more representative of the ancestral state, where the hypothesized proximal leg region is not fused to the body wall.

Towards this aim, we examined five leg gap genes, *Distalless* (*Dll*), *Sp6-9*, *dachshund* (*dac*), *extradenticle* (*exd*), and *homothorax* (*hth*), in an amphipod crustacean, *Parhyale hawaiensis*. While we have documented their expression at several developmental stages (Fig. S1), our comparative analysis does not rely solely on these expression patterns, given that expression is not always a reliable indication of function, and expression is often temporally dynamic (*5*). Instead, we have systematically knocked out these genes in *Parhyale* using CRISPR-Cas9 mutagenesis and compared this to our understanding of their function in *Drosophila* and other insects (Figs. 2, S2).

Insects have six leg segments, while *Parhyale* has seven (Fig. 1). In insects, *Dll* is required for the development of leg segments 2 – 6 (*6*-*9*). In *Parhyale*, the canonical *Dll* gene, *Dll*-e (*10*-*12*), is required for the development of leg segments 3 – 7 (Fig. 2b). In insects, *Sp6-9* (*13*) is required for the development of leg segments 1 – 6 (*14*), and in addition in *Drosophila*, loss of *Sp6-9* (i.e. D-Sp1, (*13*)) occasionally transforms the leg towards wing and lateral body wall identity (*14*). In *Parhyale*, *Sp6-9* (*13*, *15*) is required for the development of leg segments 2 – 7 (Fig. 2c), and in some legs, segment 2 is occasionally transformed towards a leg segment 1 identity (Fig S3). In *Drosophila*, *dac* is required in the trochanter through proximal tarsus (leg segments 2 – 4, and first tarsus, (*15*, *16*). *Parhyale* has two *dac* paralogs. *dac1* does not seem to be expressed in the legs or have a morphologically visible knockout phenotype. *dac2* is required to pattern leg segments 3 – 5 (Fig. 2d). *exd* and *hth* are expressed in the body wall and proximal leg segments of insects (*17*-*20*) and *Parhyale* (*21*) (Fig S1). They form heterodimers and therefore have similar phenotypes (*17*-*20*). In insects, *exd* or *hth* knockout results in deletions/fusions of the coxa through proximal tibia (leg segments 1 – 3, and proximal tibia, *17*-*20*). In *Parhyale*, *exd* or *hth* knockout results in deletions/fusions of the coxa through proximal carpus (leg segments 1 – 4, and proximal carpus; Figs. 2e, f). In both insects (*17*, *18*, *22*) and *Parhyale*, the remaining distal leg segments are sometimes transformed towards a generalized thoracic leg identity (compare Fig. 2 e, f and Fig S4). In both insects (*17*-*20*) and *Parhyale* (Fig. S4), *exd* or *hth* knockout results in deletions/fusions of body segments.

**Fig. 1.**
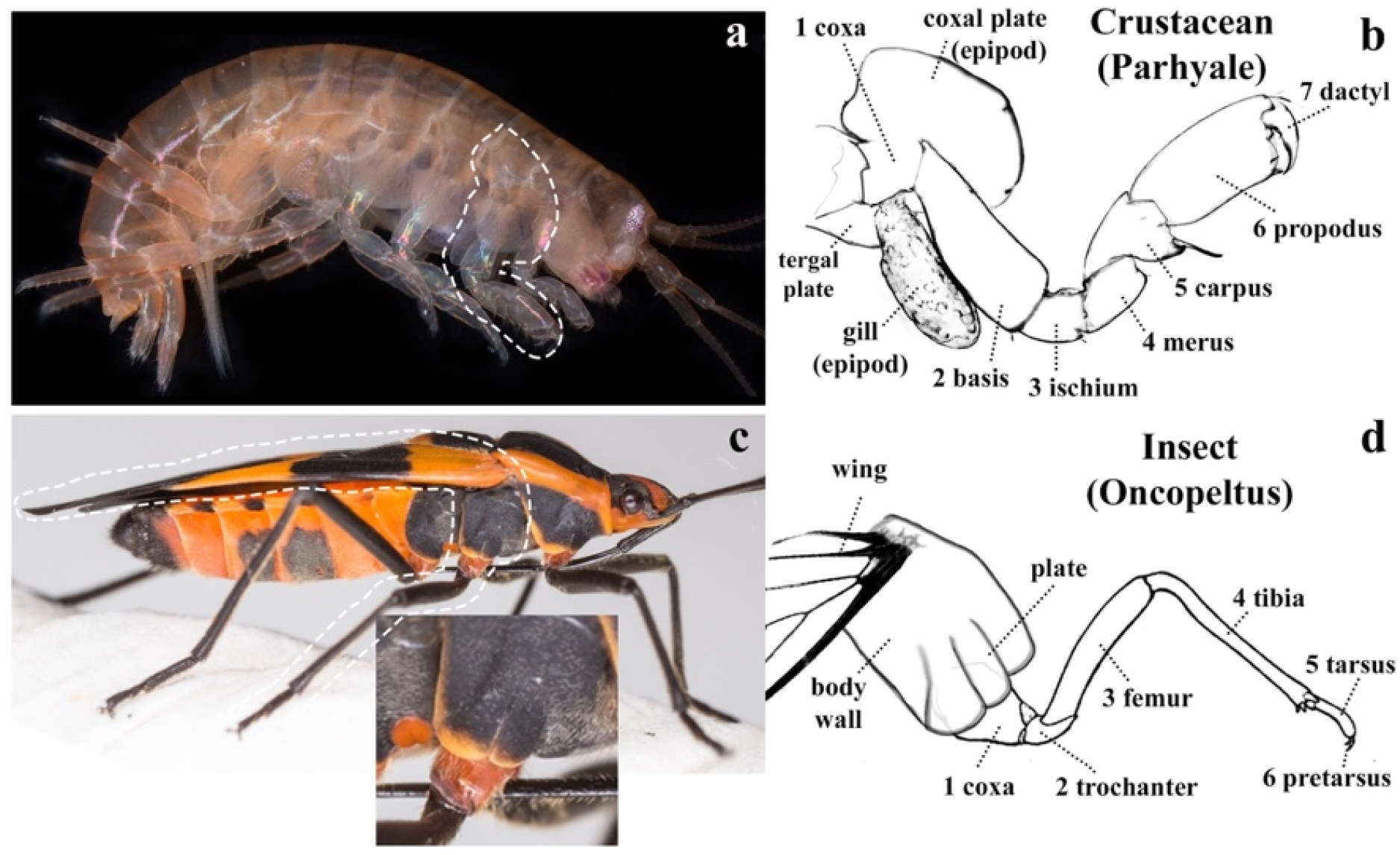
Crustacean and insect legs. (a) Adult *Parhyale*, with third thoracic leg (T3) outlined. (b) Cartoon of *Parhyale* T3. The coxal plate extends over the leg. (c) Adult *Oncopeltus*, with T2 outlined. Inset shows magnified proximal leg, with body wall plate extending over the leg. (d) Cartoon of *Oncopeltus* T2 leg.

**Fig. 2.**
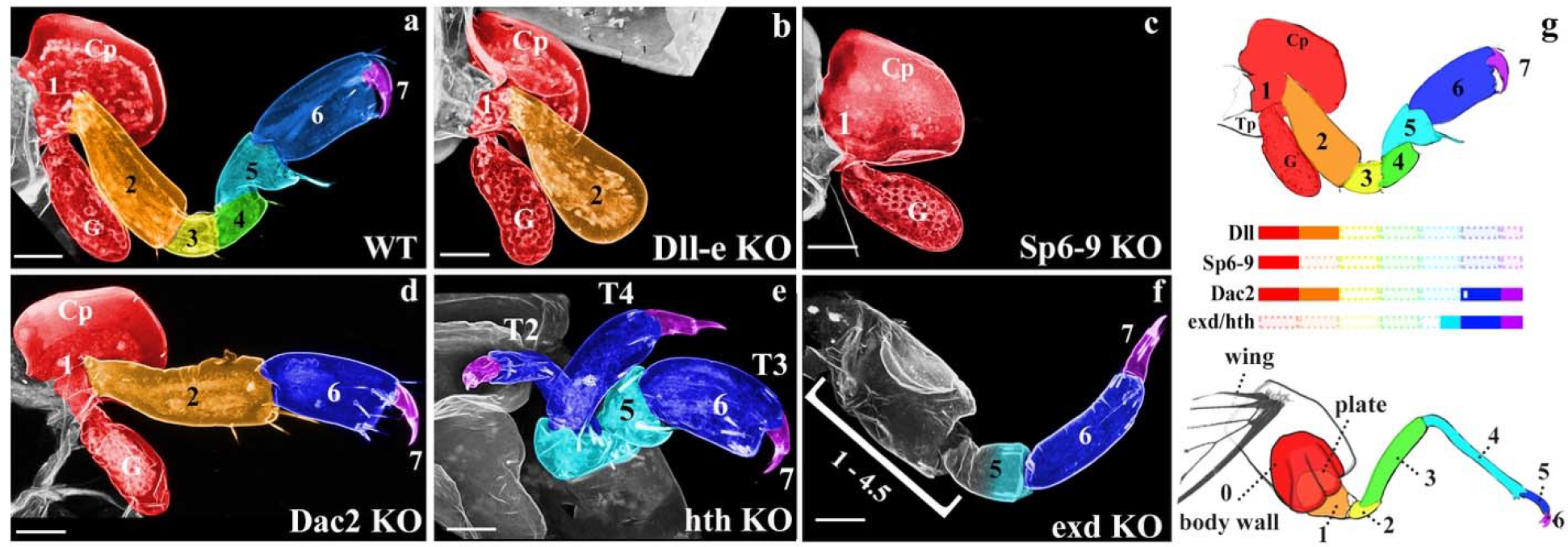
Knockout phenotypes of leg gap genes. (a-f) *Parhyale* CRISPR-Cas9 phenotypes in dissected third thoracic legs (T3). Graded cyan in f indicates deletion/fusion of proximal leg segment 5. (g) Leg gap gene function in *Parhyale* and insects aligns only if insects incorporated the red leg segment into the body wall (0). Color bars correspond to remaining leg segments following knockout, transparent bars indicate deleted leg segments. Open bar in *dac* indicates slight extension of *dac* function into tarsus 1 of insects. Coxal plate (Cp), gill (G), tergal plate (Tp). Scale bar 50um.

In summary, the expression and function of *Dll*, *Sp6-9*, *dac*, *exd*, and *hth* in *Parhyale* are shifted distally by one segment relative to insects. This shift is accounted for if insects fused an ancestral proximal leg segment into the body wall (Fig. 2g). Thus, there is a one-to-one homology between insect and *Parhyale* legs, displaced by one segment, such that the insect coxa is homologous to the crustacean basis (*23*), the insect femur is the crustacean ischium, and so on for all leg segments. This also means that part of the insect body wall is homologous to the crustacean coxa.

Clark-Hachtel (accompanying manuscript) show that the plates on the *Parhyale* basis, coxa, and lateral body wall are epipods. The body wall epipod is notable, because epipods are characteristic of leg segments (*23*). This suggested to us that part of the *Parhyale* body wall might actually be an additional leg segment. In fact, most groups of crustaceans have an additional proximal leg segment, the precoxa (Fig. 3a). To determine whether *Parhyale* retains the precoxa, we examined dissected *Parhyale* using confocal and brightfield microscopy. We identified a precoxal structure that meets the criteria for a true leg segment: it protrudes conspicuously from the body wall; it forms a true, muscled joint; and it extends musculature to another leg segment (Figs. 3 and S5, (*23*-*25*)). Importantly, the plate does not emerge from the body wall, but from the precoxa (Fig. 3e), like the plates of the coxa and basis. Thus, much of what appears to be lateral body wall in *Parhyale* is in fact proximal leg.

**Fig. 3.**
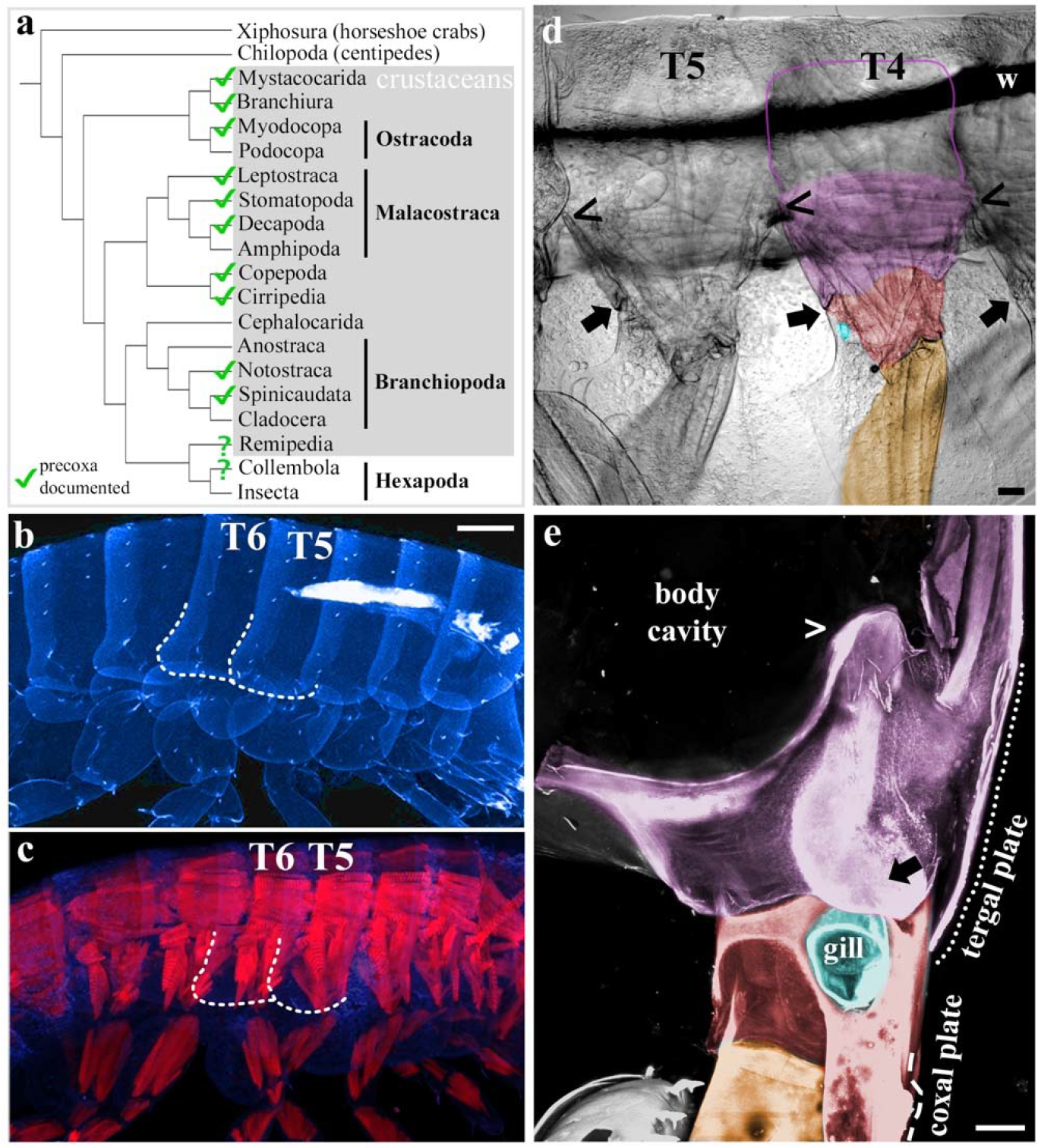
Evidence for a precoxa in *Parhyale*. (a) Phylogeny based on Oakley 2012, precoxa references in supplements. (b) Confocal image of *Parhyale* hatchling, autofluorescent cuticle in blue. T5, T6 tergal plates (dotted outlines). (c) Confocal image of *Parhyale* hatchling, autofluorescent cuticle in blue, muscle phalloidin stain in red. Compare blocks of simple, anterior-posterior muscles of the body to orthogonal, complexly arranged muscles of the leg segments. Note overlap between tergal plate (dotted lines) and orthogonal leg muscle. (d) Brightfield image of right half of *Parhyale,* sagittal dissection, internal tissues removed, lateral view. Wire used to position sample (w). The same orthogonal muscles in b are visible as striations extending above the wire. The precoxa forms a joint with the coxa (*47*) (arrow). The dorsal limit of the precoxa is unclear: a conservative estimate is to begin at the joint (arrow) and follow the leg up to where it meets the adjacent leg, denoted by (<); however, the orthogonal muscle striations continue farther up (pink outline). Either way, the precoxa protrudes quite a bit from the body wall. (e) Posterior-lateral view of right T6, looking edge-on at tergal plate. The tergal plate (dotted outline) emerges from the precoxa (contiguous pink between ←, >, and ---), just as the coxal plate (dashed line) emerges from the coxa. In c, d, coxa is red (coxal plate not shaded), gills (teal) partially cut for visibility, basis orange, precoxa pink. Note that all three plates (tergal, coxal, and basal) form contiguous cuticle with their leg segment, i.e. there is no distinguishing suture. Scale bar 100um.

If the insect coxa is homologous to the crustacean basis, what happened to the leg segments corresponding to the ancestral crustacean precoxa and crustacean coxa in insects? If these two leg segments became incorporated into the body wall, then one would expect to find two leg segments and two epipods dorsal to the insect coxa (Fig. 4a). As predicted, two leg-like segments can be observed proximal to the coxa in basal hexapods (*1*) including collembolans (*26*), as well as in the embryos of many insects (*27*-*29*), where these two leg-like segments flatten out before hatching to form the lateral body wall (*1*, *26*-*31*). Insects also appear to have two epipods dorsal to the insect coxa, because when “wing” genes are depleted in insects via RNAi, two distinct outgrowths are affected, the wing and the plate adjacent to the leg (Fig. 1c, (*32*-*35*)).

**Fig. 4.**
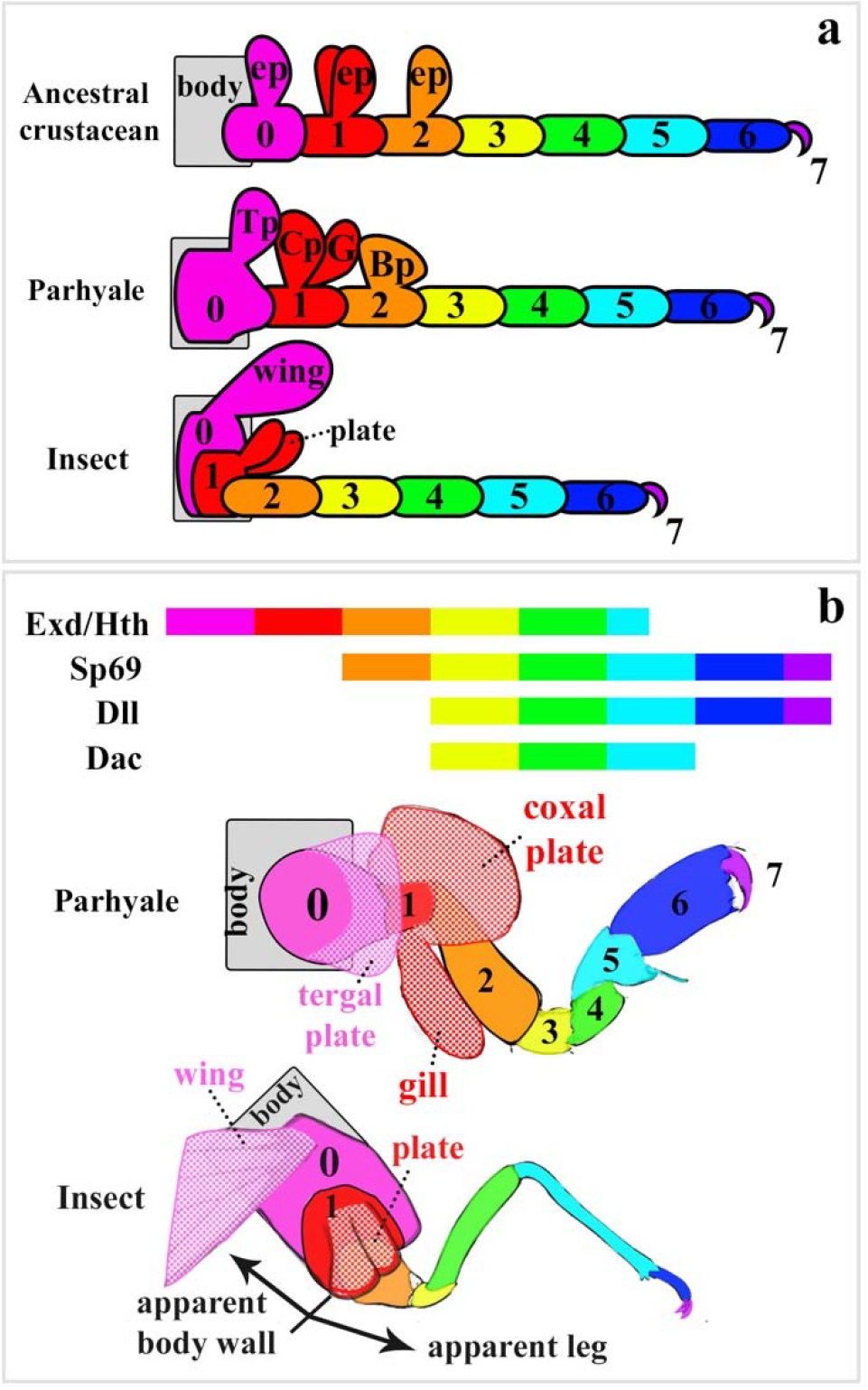
Proposed leg segment homologies (colors) between insects, *Parhyale*, and an ancestral crustacean (a) based on gene function alignment (b). Ancestral precoxa epipod (pink ep), *Parhyale* tergal plate (Tp), and insect wing are homologous (pink). Ancestral coxa epipod, *Parhyale* coxal plate (Cp) and gill (G), and insect plate (see Fig. 1c) are homologous (red). *Parhyale* basal plate (Bp). Insect numbering based on crustaceans.

Based on these data, insects incorporated two ancestral leg segments, the precoxa and crustacean coxa, into the body wall (Fig. 4a). Thus, like *Parhyale*, much of what appears to be lateral body wall in insects is in fact proximal leg. Clark-Hachtel’s interpretation of the dual origin theory proposes that these two leg segments and their two epipods fused to form the wing. While we agree that both leg segments may contribute wing muscle, we propose that only the more dorsal precoxa epipod formed the insect wing, while the more ventral crustacean coxa epipod formed the insect plate (Fig. 4b).

Our results may settle the long-standing debate regarding the origin of insect wings as derived from either an epipod of the leg or from body wall. Our model accounts for all observations in favor of either of these hypotheses, including the dorsal position of insect wings relative to their legs, the loss of ancestral leg segments in insects, the two-segmented morphology of the insect subcoxa in both embryos and adults, the complex musculature for flight, and the shared gene expression between wings and epipods. Our model also explains the apparent “dual origin” of insect wings from both body wall and leg epipod: much of what appears to be insect body wall is in fact the remnant of two ancestral leg segments and their epipods.

In fact, a number leg-associated outgrowths in arthropods could be explained by this model, in addition to insect wings. The Daphnia carapace(*36*) is the epipod of the precoxa(*37*); the *Oncopeltus* small plate outgrowth (Fig. 1c) is the epipod of the crustacean coxa; and the thoracic stylus of jumping bristletails (Fig. 4, st) is the epipod of the crustacean basis(*38*, *39*). This also explains many insect abdominal appendages, like gills(*40*), gin traps(*33*), prolegs(*41*), and sepsid fly appendages(*42*), which are often proposed as de novo structures(*43*-*45*). However, most insects form abdominal appendages as embryos(*40*, *46*), some even with an epipod nub, but these fuse to the body wall before hatching to form the sternites(*28*, *39*). The existence of insect abdominal appendages is supported by a re-analysis of the expression of *Sp6-9* and its paralog, *buttonhead*, in insect embryos in a previous study (*13*). According to the leg segment homology model presented here (Fig. 4), the paired dots of *btd* expression on each abdominal segment of insect embryos demonstrates that these appendages are comprised of three leg segments: the precoxa (pink), crustacean coxa (red), and insect coxa (orange). These abdominal appendages are truncated, lacking all distal appendages from the trochanter (yellow) down, because *Dll* and *dac*, which mark the trochanter and more distal leg segments, are not expressed in the insect abdomen. Thus, rather than de novo co-options, abdominal appendages were always there, persisting in a truncated, highly modified state, and de-repressed in various lineages to form apparently novel structures. This provides a model for how insect wings can be both homologous to the epipod of the crustacean precoxa, and yet not be continuously present in the fossil record: epipod fields may persist in a truncated state, perhaps only visible as a nub in the embryo. We therefore propose cryptic persistence via truncation as a general mechanism for the origin of apparently novel structures that appear to be derived from serial homologs, rather than the current model of extensive gene co-option.

## Acknowledgments

We thank Courtney Clark-Hachtel and Yoshinori Tomoyasu for sharing their preprint.

## Funding

This work is supported by the National Science Foundation (NSF) (to NHP), NSF Graduate Research Fellowship (to HSB).

## Author contributions

H.S.B. and N.H.P. conceived of the experiments. H.S.B. performed the experiments, conceived of model, and wrote the manuscript. N.H.P. edited and revised the manuscript.

## Competing interests

Authors declare no competing interests.

## Data and materials availability

All data is available in the main text or the supplementary materials.

